# DeepMosaic: Control-independent mosaic single nucleotide variant detection using deep convolutional neural networks

**DOI:** 10.1101/2020.11.14.382473

**Authors:** Xiaoxu Yang, Xin Xu, Martin W. Breuss, Danny Antaki, Laurel L. Ball, Changuk Chung, Chen Li, Renee D. George, Yifan Wang, Taejeoing Bae, Alexej Abyzov, Liping Wei, Jonathan Sebat, NIMH Brain Somatic Mosaicism Network, Joseph G. Gleeson

## Abstract

Mosaic variants (MVs) reflect mutagenic processes during embryonic development^1^ and environmental exposure^2^, accumulate with aging, and underlie diseases such as cancer and autism^3^. The detection of MVs has been computationally challenging due to sparse representation in non-clonally expanded tissues. While heuristic filters and tools trained on clonally expanded MVs with high allelic fractions are proposed, they show relatively lower sensitivity and more false discoveries^4–9^. Here we present DeepMosaic, combining an image-based visualization module for single nucleotide MVs, and a convolutional neural networks-based classification module for control-independent MV detection. DeepMosaic achieved higher accuracy compared with existing methods on biological and simulated sequencing data, with a 96.34% (158/164) experimental validation rate. Of 932 mosaic variants detected by DeepMosaic in 16 whole genome sequenced samples, 21.89-58.58% (204/932-546/932) MVs were overlooked by other methods. Thus, DeepMosaic represents a highly accurate MV classifier that can be implemented as an alternative or complement to existing methods.

---

Postzygotic mosaicism describes a phenomenon whereby cells arising from one zygote harbor distinguishing genomic variants^1, 10^. MVs can act as recorders of embryonic development, cellular lineage, and environmental exposure. They accumulate with aging, play important roles in human cancer progression^3, 10^, and are implicated in over 200 non-cancerous disorders^11, 12^. Collectively, estimates are that MVs contribute to 5-10% of the ‘missing genetic heritability’ in more than 100 human disorders^11, 13^.

Compared with the higher allelic fractions (AF) of 5-10% found in clonal tumors or pre-cancerous mosaic conditions, AFs found in non-clonal disorders, or neutral variants used for lineage studies, are typically present at much lower AFs. Existing methods, however, based on classic statistical models like MuTect2^9^ and Strelka2^7^ and heuristic filters are often optimized for the high fraction variants in cancer with relatively high variant AFs. Similarly, because of their conceptual origin in cancer, most existing programs including the more recent NeuSomatic^14^, also require matched control samples. This can be problematic when mutations are present across different tissues (‘tissue shared’ mosaicism), or when only one sample is available.

Newer methods that aim to overcome these limitations, such as MosaicHunter^5^ or MosaicForecast^4^, are based conceptually similarly on the use of features extracted from raw data, rather than the sequence and alignment themselves, or replace the filters with traditional machine-learning methods. While these are a useful proxy, they only represent a limited window into the sheer wealth of information. Because of these limitations, researchers often resort to visual inspection of raw sequence alignment in a genome browser, a so-called ‘pileup’, to distinguish artifacts from true positive variants^15^. This is a laborious and low-throughput process that allows spot checking, but cannot be implemented on a large scale for variant lists numbering in the thousands for programs like MuTect2.

Image-based representation of pileups and the application of deep convolutional neural networks represents a potential solution for these limitations. Previous attempts like DeepVariant^14^ were successful in detecting heterozygous or alternative homozygous single nucleotide variants (SNVs) from direct representation of aligned reads by using deep neural networks. The DeepVariant genotype model, unfortunately, did not consider a mosaic genotype, and lacked orthogonal validation experiments. Here we introduce DeepMosaic comprising two modules: a visualization module for image-based representation of SNVs, which forms the basic input for a convolutional neural network (CNN)-based classification module for mosaic variant detection. Seven different biological and computationally simulated dataset as well as amplicon validation were used to train and benchmark DeepMosaic.

To automatically generate a useful visual representation similar to a browser snapshot, we developed the visualization module of DeepMosaic (DeepMosaic-VM, Fig.1a-d). The input for this visualization is short-read WGS data, processed with a GATK current best-practice pipeline (insertion/deletion, or INDEL, realignment and base quality recalibration). DeepMosaic-VM processes this input into an ‘RGB’ image, representing a pileup at each genomic position. In contrast to a regular browser snapshot, we encode sequences as different intensities within one channel, and use other channels for base quality and strand orientation. We further split the pileup of reference reads and alternative reads based on the reference genome information (Fig. 1a-d), to improve visualization and allow assessment of mosaicism at a glance.

**Fig. 1.**
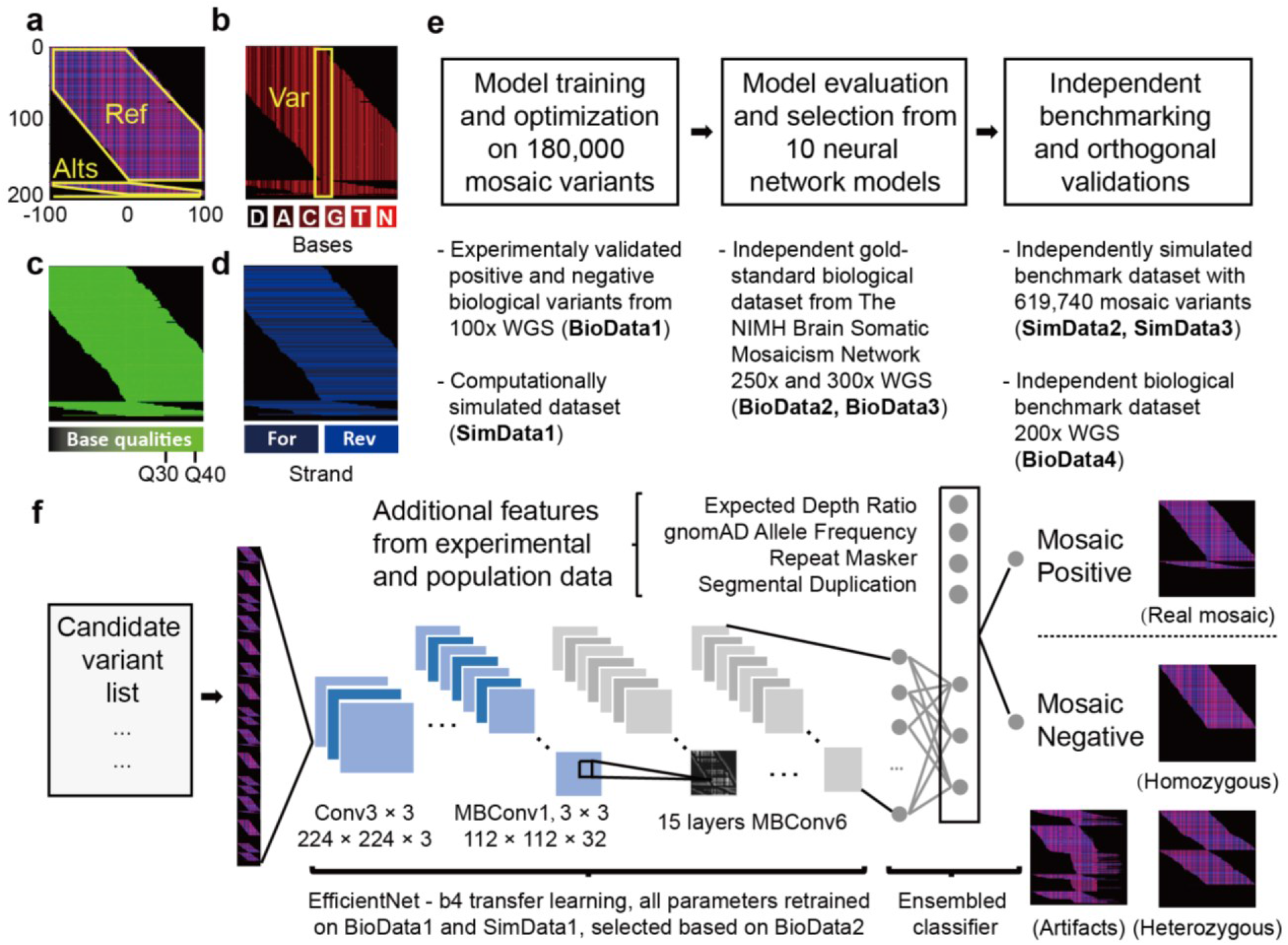
Image representation, model training strategies, and framework of DeepMosaic. **a**, DeepMosaic-VM: Composite RGB image representation of sequenced reads separated into “Ref” - reads supporting the reference allele; or “Alts” - reads supporting alternative alleles; each outlined in yellow. **b**, Red channel of the compound image contains base information from BAM file. “D” - deletion; “A” – Adenine; “C” – cytosine; “G” – guanine; “T” – thymine; “N” – low-quality base. Yellow box: Var: candidate position, centered in the image. **c**, Green channel: base quality information. Note that channel intensity was modulated in this example for better visualization. **d**, Blue channel: strand information (i.e. forward or reverse). **e**, Model training, model selection, and overall benchmark strategy for DeepMosaic-CM (Methods and Supplementary Fig. 1). Ten different convolutional neural network models were trained on 180,000 experimentally validated positive and negative biological variants from 29 WGS data from 6 individuals sequenced at 100x^16, 17^ (BioData1), as well as simulated data with different AFs (SimData1) resampled to a different depth. Models were evaluated based upon an independent gold-standard biological dataset from the 250x WGS data of the Reference Tissue Project of the Brain Somatic Mosaicism Network^23^ (BioData2) as well as an independent 300x WGS dataset from the Brain Somatic Mosaicism Network Capstone project^18^ (Biodata3). DeepMosaic was further benchmarked on 16 independent biological datasets from 200x WGS data^27^ (BioData4) as well as 619,740 independently simulated variants (SimData2 and SimData3). Deep amplicon sequencing was carried out as an independent evaluation on variants detected by different software (Supplement Table 1). **f**, Application of DeepMosaic-CM in practice. Input images are generated from the candidate variants. 16 convolutional layers extracted information from input images. Population genomic features were assembled for final output. Images of positive and negative variants are shown as examples. Conv: convolutional layers; MBConv: mobile convolutional layers.

The classification module of DeepMosaic (DeepMosaic-CM) is a CNN-based classifier for MVs. We trained 10 different CNN models with more than 180,000 image-based representations from both true-positive and true-negative biological variants in several recently published high-quality experimentally validated public datasets^16–18^, and computationally simulated reads with added MVs (employing Illumina HiSeq error models) across a range of AFs and depths (Fig. 1e, Methods and Supplementary Fig. 1a-b) to select a model with optimal performance. To ensure its resemblance of real data, we controlled the distribution of AFs in the training set (Supplementary Fig. 1c). In addition, a range of expected technical artifacts, including multiple alternative alleles, homopolymers, and alignment artifacts, were manually curated and labeled negative in the training set to represent expected pitfalls that often result in false positive mosaic calls for other programs (Supplementary Fig. 1d).

To further expand training across a range of different read depths, the biological training data were also up- and down-sampled to obtain data at read depths ranging from 30x to 500x (Supplementary Fig. 1e), which includes the most commonly used depths for WGS in current clinical and scientific settings. In addition to the output from DeepMosaic-VM, we further incorporated population genomic and sequence features (e.g. population allele frequency, genomic complexity, ratio of read depth), which are not easily represented in an image, as input for the classifier (Fig. 1f). Depth ratios were calculated from the expected depth and used to exclude false positive detections from potential copy number variations (CNVs). gnomAD population allelic frequencies were used to exclude common variants. Segmental duplication and repeat masker regions were used to exclude 24% of low complexity regions genome-wide.

Ten different CNN architectures were trained on 180,000 training variants described above. The CNN models included Inception-v3^19^, which was used in DeepVariant; Deep Residual Network^20^ (Resnet) which was used in the control-dependent caller NeuSomatic; Densenet^21^ and 7 different builds of EfficientNet^22^, for its high performance on rapid image classification (Methods, Supplementary Fig. 2a). Each model was trained on the data described above with 5 to 15 epochs to optimize the hyper-parameters until training accuracies plateaued (>0.90).

To compare the different models after training and to contrast models trained with distinct datasets, we employed an independent gold-standard validation dataset of ~400 MVs from one brain sample^23^ (BioData2, Methods) and another amplicon-validated dataset from 18 samples from one individual^18^ (BioData3, Methods). On these, EfficientNet-b4 showed the highest accuracy, Matthews’s correlation coefficient, and true positive rate when trained for 6 epochs (Supplementary Fig. 2b). We thus selected this model as the default model of DeepMosaic-CM (Supplementary Fig. 3a and Fig. 1f). Additional EfficientNet-b4 models trained on the 1:1 mixture of biological data and simulated data showed similar performance compared with biological data only training set but much higher specificity compared with models trained only on simulated data (Supplementary Fig. 2c).

To uncover the information prioritized by the selected default model, we used a gradient visualization technique with guided backpropagation^24^ to highlight the pixels with guiding classification decisions (Supplementary Fig. 3b). The results suggested that the algorithm not only recognized the edges for reference and alternative alleles, but also integrated additional available information, such as insertion/deletions in the sequences, overall base qualities, alignment artifacts, and other features which may not be extracted by digested feature-based methods.

We evaluated the performance of DeepMosaic using 20,265 variants from the above training data that were hidden from model training and selection. The receiver operating characteristic (ROC) curve and precision-recall curves on the hidden validation dataset showed >0.99 area under the curve for a range of coverages (30x ~500x, Supplementary Fig. 4a and 4b) across a range of AFs (Supplementary Fig. 4c and 4d), demonstrating high sensitivity and specificity.

Next, we benchmarked DeepMosaic’s performance relative to other detection software, using data generated from two distinct sequencing error models to test for its utility on general sequencing data. We compared the performance of DeepMosaic with the widely used MuTect2 (paired mode), Strelka2 (somatic mode) with heuristic filters, MosaicHunter (single mode), and MosaicForecast (Methods). We generated two additional computationally simulated datasets of 439,200 and 180,540 positions based on the error model of a different Illumina sequencer with similar methods as the training set (NovaSeq, SimData2, Methods) or a similar ratio of true positive and true negative labels as real biological data^18^ by replacing reads from the ‘Genome in a Bottle’ sample HG002 (NA24345, SimData3, Methods)^25, 26^, with AF ranges from 1% to 25%, and depth ranges from 50x to 500x. MuTect2 paired methods and Strelka2 somatic mode used simulated mutated samples as “tumor” and simulated reference or original HG002 samples as “normal” for their paired modes. DeepMosaic showed equal or better performance than all other methods tested, especially for low allelic fraction variants (Fig. 2 and Supplementary Fig. 5), noticeably, even for low read depth data; and it performed better than methods that have additional information from paired samples. Overall DeepMosaic showed a 1.5-3 fold increase of the detection sensitivity for AFs under 3% compared with other methods (Fig. 2b), with comparable specificity (Fig 2a). This is likely because our models integrate additional genomic sequence and quality information from the original BAM file (Supplementary Fig. 3b), and are capable of distinguishing mosaic variants from different Illumina error models.

**Fig. 2.**
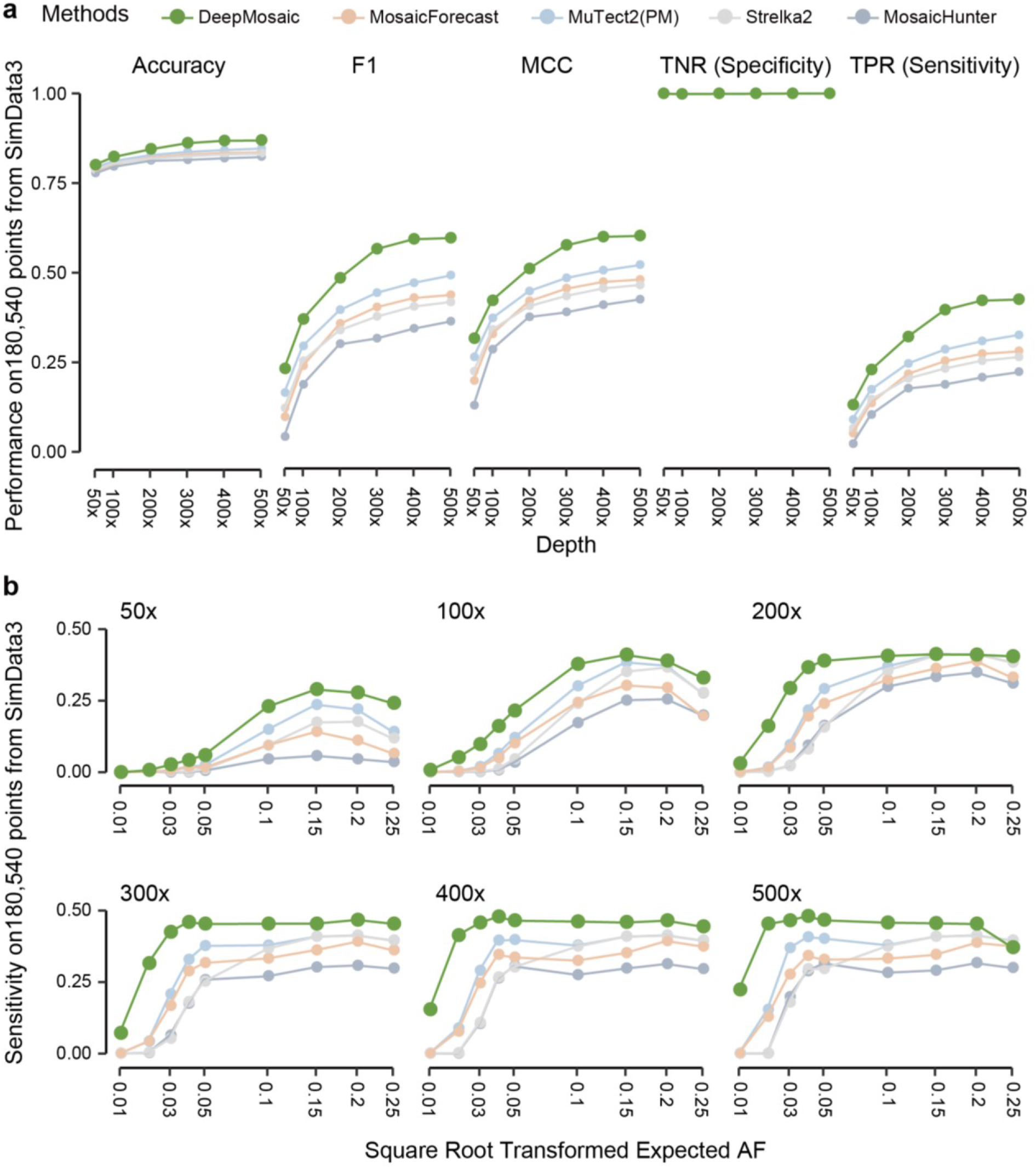
DeepMosaic performance on simulated benchmark variants. **a,** Benchmark test on 180,540 genomic positions (SimData3) generated by replacing reads from biological data with simulated variants. DeepMosaic showed higher accuracy, F1 score, MCC (Matthews correlation coefficient), sensitivity, and comparable specificity compared with widely accepted methods for mosaic variant detection. **b,** Sensitivity of DeepMosaic and other mosaic callers on SimData3 at simulated read depths and AFs. DeepMosaic performed equally well or better than other tested methods, especially at lower read-depths and lower expected AFs.

To exclude limitations resulting from benchmarking with simulated data and demonstrate that models trained on PCR-amplified libraries are also useful for PCR-free sequencing libraries, we extended benchmarking to biological data. We performed the same comparison on our recently published 200x WGS dataset^12^ with 16 samples (blood and sperm) from 8 healthy individuals^27^. Paired methods compared two samples from the same individual, and control-independent samples used a published dataset of a panel of normals^28^. Variants detected by MuTect2 (paired mode), Strelka2 (somatic mode) and MosaicHunter (single mode) were subjected to a series of published heuristic filters^27, 28^. As we had access to the biological samples, we also performed orthogonal validation, using deep amplicon sequencing of 241 randomly selected MVs with a representative AF distribution compared to the complete candidate variant list (Methods, Fig. 3a and 3b, Supplementary Table 1).

**Fig. 3.**
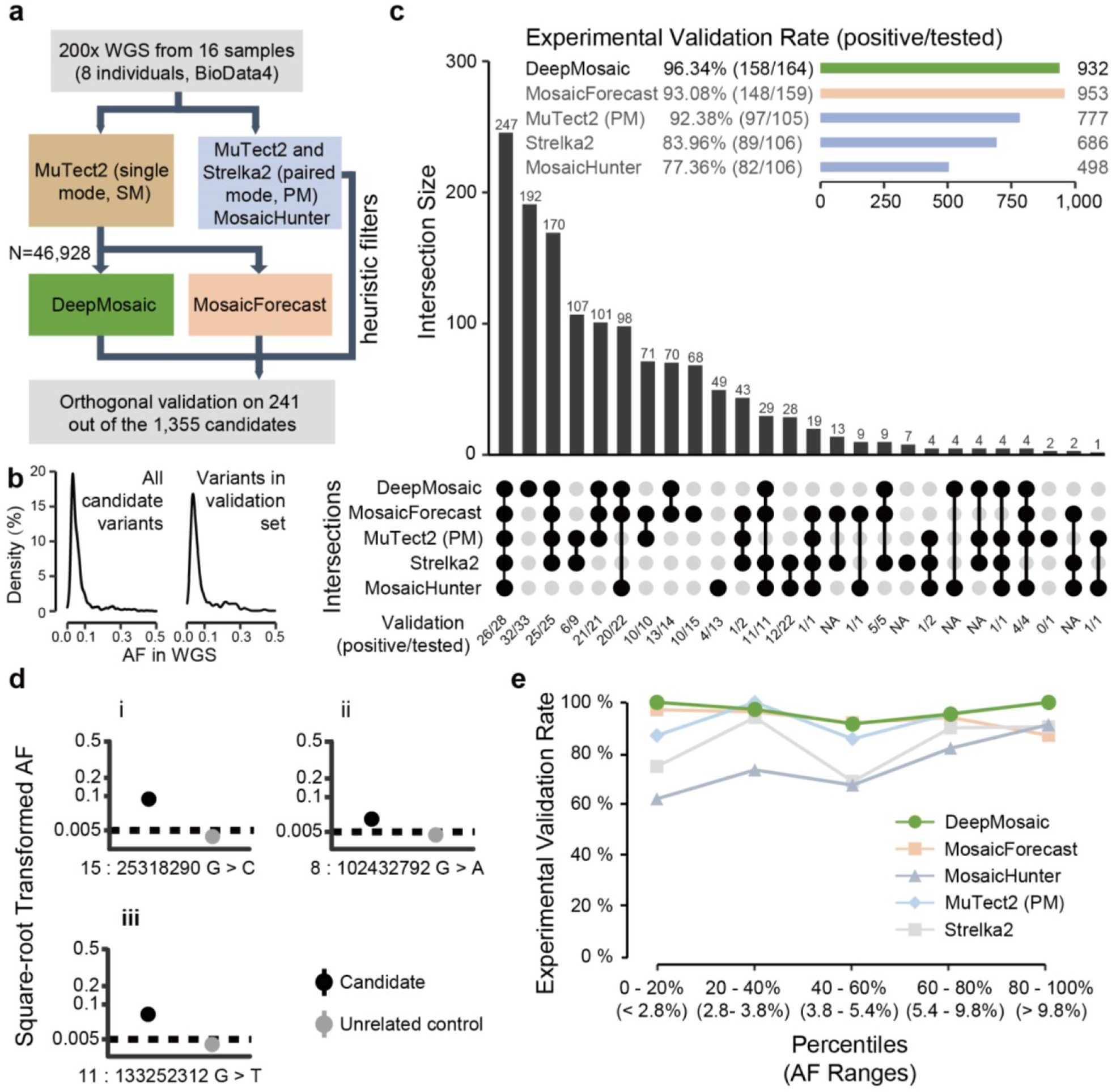
DeepMosaic performance validated on biological data. **a**, DeepMosaic and other mosaic variant detection methods were applied to 200x whole-genome sequencing data from 16 samples, which were not used in the training or validation stage for any of the listed methods (BioData4). Raw variant lists were either obtained by comparing samples using a panel-of-normal^28^ strategy with MuTect2 single mode, between different samples from a same individual using MuTect2 paired mode or Strelka2 somatic mode, or detected directly without control with MosaicHunter single mode with heuristic filters^27^. A total of 46,928 candidate variants from MuTect2 single mode were analyzed by DeepMosaic and MosaicForecast. Orthogonal validation with deep amplicon sequencing was carried out on a total of 241 variants out of the 1355 candidates called by at least one method. **b**, Distribution of AFs of the whole candidate mosaic variant list and the 241 randomly selected variants. **c**, Comparison of validation results between different mosaic variant calling methods, ‘UpSet’ plot shows the intersection of different mosaic detection methods and the validation result of each category. Variants identified by DeepMosaic showed the highest validation rate on biological data. **d**, Examples of validated variants called by DeepMosaic and MosaicForecast (i), only by DeepMosaic (ii), or by DeepMosaic and other traditional methods (iii). **e**, Comparison of validation rate in different AF range percentage bins of variants. DeepMosaic showed the highest validation rate at a range of AFs, approximately 48 experimentally validated variants are shown in each AF bin.

As expected from the test of the computationally generated data, DeepMosaic showed the highest overall validation rate (96.34%, 158/164) among all 5 methods (Fig. 3c), demonstrating the power of DeepMosaic that models trained on PCR-amplified biological data and simulated data can accurately classify these PCR-free biological data. Of the 932 MVs detectable by DeepMosaic, 21.89% (204/932, 33/34 experimentally validated) were overlooked by MosaicForecast, 58.58% (546/932, 96/98 validated) overlooked by MosaicHunter, 50.32% (469/932, 90/94 validated) overlooked by Strelka2 (somatic mode) with heuristic filters, 43.13% (402/932, 81/85 validated) overlooked by MuTect2 (paired mode) with heuristic filters^27^. DeepMosaic also accurately detected variants with relatively low AFs (Fig. 3d). Finally, DeepMosaic outperformed other methods across most of the AF bins (Fig. 3e).

In current practice, researchers often combine multiple programs in one variant detection pipeline to detect different categories of MVs^27–29^. We thus further compared DeepMosaic with different pipelines used in recent publications, using data from 200x WGS of the 16 samples^27^: 1] With the MosaicForecast pipeline^4^, which uses MuTect2 single mode (each sample compared with the publicly available panel of normal) as input; 2] With what we call the M2S2MH pipeline, which we recently published^27^, combining MuTect2 paired mode (i.e. compared between different samples from a same individual), Strelka2 somatic mode and MosaicHunter single mode followed by a series of heuristic filters (Supplementary Fig. 6a). Of the 932 MVs identified by DeepMosaic, 78.11% (728/932, 125/130 validated) overlapped with MosaicForecast and 60.09% (560/932, 87/91 validated) overlapped with M2S2MH. In contrast, 21.89% (204/932, 33/34 validated) were undetected by MosaicForecast, and 39.91% (372/932, 71/73 validated) were overlooked by M2S2MH. These variants uniquely detected by DeepMosaic all showed validation rate > 97% (Supplementary Fig. 6b and 6c), demonstrating that DeepMosaic can accurately detect a considerable number of variants undetectable by widely used methods.

To test the performance of these samples on data widely curated clinically, we compared detection sensitivity for genome samples with standard WGS read depth, by down-sampling blood-derived data from a 70-year old healthy individual, in whose blood we observed the highest number of mosaic variants (due to clonal hematopoiesis^27^). As all programs had high validation on this sample at 200x, the recovery rate was used to distinguish the ability of different programs to detect clonal hematopoiesis variants. DeepMosaic showed similar recovery in the down-sampled data (Supplementary Fig. 7) as M2S2MH and slightly outperformed MosaicForecast at 100x and 150x. We found that the performance of DeepMosaic was not substantially influenced by the read depth according to the down-sampling benchmark on biological data.

To understand whether different pipelines had unique strengths or weaknesses, we separated all the detected variants into 7 groups (G1-G7) based upon sharing between different pipelines, Supplementary Fig. 7a). DeepMosaic specific variants showed similar base substitution features compared with other methods (Supplementary Fig. 7b). Similar to the computationally derived data, we found that DeepMosaic recovered additional low AF MVs with high accuracy (validation rate 95%, Supplementary Fig. 7c). Finally, we summarized the genomic features of variants detected by DeepMosaic and other pipelines. All caller groups report similar ratios of intergenic and intronic variants (Supplementary Fig. 8a). Analysis of other genomic features showed DeepMosaic-specific variants (G1) expressed consistency with other groups (Supplementary Fig. 8b), reflecting that the low-fraction variants detectable only by DeepMosaic do not represent technical artifacts.

While we propose DeepMosaic as a powerful tool for mosaic variant detection, it currently is underpowered for mosaic INDELs and mosaic repetitive variant detection which might be error-prone in the genome. In practice, MosaicForecast can detect mosaic INDEL variants with high accuracy, while M2S2MH has good performance for tissue-specific variants due to the inclusion of additional information from the “normal” comparison sample. Thus different methods complement one another.

DeepMosaic is the first image-based tool for the accurate detection of mosaic SNVs from short-read sequencing data and does not require a matched control sample. Compared with NeuSomatic that compresses all the bases in a genomic position into 10 features^6^, DeepMosaic-VM provides complete representation of information present in the BAM file. Compared with other re-coding methods like DeepVariant^14^, DeepMosaic-CM has the ability to define MVs as an independent genotype and DeepMosaic-VM can be applied as an independent variant visualization tool for the user’s convenience. To further integrate population information not present in the raw BAM, 4 different features are also integrated in DeepMosaic to facilitate classification.

Despite the unique features from image representation and a neural network based variant classifier, DeepMosaic can reproducibly identify the majority (~70%) of variants detectable by conventional methods; in addition, however, this unique architecture results in higher sensitivity, and the detection of variants with relatively lower AF both in simulated and experimentally derived data validated by orthogonal experiments. DeepMosaic shows a drop of sensitivity at higher AF, likely due to our inclusion of depth ratio which help to avoid false-positive calls from CNV. The higher sensitivity at lower AFs will make it a good complement for other methods.

Both down-sampled biological data in blood of an individual with advanced age and computationally generated data showed that DeepMosaic has the potential to identify variants at relatively high sensitivity and high accuracy for WGS at depths as low as 50x. Clonal hematopoiesis in blood without a known driver mutation is reported^30^, but can be difficult to detect because of technical limitations induced by noise and lower supporting read counts^31^. For the past 10-15 years, hundreds of thousands of whole-genome sequencing datasets from clinical, commercial, or research labs have been generated at relatively low depth, but most have not been subjected to unbiased mosaicism detection due to lack of sufficiently sensitive methods. DeepMosaic could enable a genome-level unbiased detection of mutations that requires only conventional sequencing data.

By using a training set comprising representative technical artifacts such as homopolymers and truncated reads, DeepMosaic acquired the power to distinguish biologically true positive variants from false positives, which might be filtered out directly by rule-based methods like MosaicHunter^5^ or MosaicForecast^4^. We demonstrated that training the models on a mixture of ~1:1 simulated and biological data does not adversely affect performance on an independent biological evaluation set. We also demonstrated that DeepMosaic works well for various Illumina short read sequencing platforms applying different library preparation strategies (PCR-amplified and PCR-free).

Although the EfficientNet-b4 performed best, we provide all pre-trained models (Densenet, EfficientNet, Inception-v3, and Resnet) as DeepMosaic-CM modules on GitHub. Users are provided with the options to prepare their own data with labeled genotypes for model training for DeepMosaic, to generate data-specific, personalized models, to test other potential factors influencing detection sensitivity such as the ratio of positive:negative labels, and to increase the detection specificity for DeepMosaic on specialized data sets. For instance, homopolymers and tandem repeats are increasingly recognized as important in disease and development, but are currently not detected with DeepMosaic, because of the limited training data; however, users with specialized data sets could remedy this problem by re-training.

Likewise, gnomAD population AF features used in this study also rely on a matched ancestry background to avoid population stratification. Annotations such as gene names, variant functional annotations, gnomAD allelic frequency, homopolymer and dinucleotide repeat annotation, as well as segmental duplication and UCSC repeat masker regions are provided in the final output to facilitate customization, as described at the GitHub homepage of DeepMosaic (https://github.com/Virginiaxu/DeepMosaic). Finally, apart from MuTect2 single mode, DeepMosaic can also process variant lists generated by multiple methods such as the GATK HaplotypeCaller with ploidy 50 or 100^23^. Thus, DeepMosaic can be used directly as is, or can be customized to the needs of the end users, providing an adaptable and efficient mosaic variant detection workflow.

## Methods

### Curation of training and benchmark data

#### SimData1

For the initial training procedure, 10,000 variants were randomly generated on chromosome 22 to get the list of alternative bases. Pysim^32^ was then used to simulate paired-end sequencing reads with random errors generated from the Illumina HiSeq sequencer error model. Alternative reads were generated by replacing the genomic bases with the alternative bases in the list, with the same error model. Alternative and reference reads were randomly mixed to generate an alternative AF of 0, 1, 2, 3, 4, 5, 10, 15, 20, 25, and 50%. The data were randomly sampled for a targeted depth of 30, 50, 100, 120, 150, 200, 250, 300, 400, and 500x. FASTQ files were aligned to the GRCh37d5 human reference genome with BWA (v0.7.17) *mem* command. Aligned data were processed by GATK (v3.8.1) and Picard (v2.18.27) for marking duplicate, sorting, INDEL realignment, base quality recalibration, and germline variant calling. The up- and down-sampling expanded this dataset into a pool of 990,000 different variants. Depth ratios were calculated as defined. To avoid the situation that randomly generated mutations falls on a common SNP position in the genome, which would bias the training and benchmarking, gnomAD allele frequencies were randomly assigned from 0 to 0.001 for simulated mosaic positive and from 0 to 1 for simulated negative variants, which were established as homozygous or heterozygous.

#### BioData1

Variant information and raw sequencing reads from 80-120x PCR-amplified PE-150 WGS data of 29 samples from 6 normal individuals were extracted from published data^16, 17^ on SRA (SRP028833, SRP100797, and SRP136305). 921 variants identified from WGS of samples from different organs of the donors and validated by orthogonal experiments were selected and labeled as mosaic positive. 492 genomic positions from the control samples validated with 0% AF were selected and labeled as negative. 162 variants with known sequencing artifacts were first filtered by MosaicHunter, manually selected and labeled as negative. The 1575 genomic positions were also down-sampled and up-sampled for a targeted depth of 30, 50, 100, 150, 200, 250, 300, 400, and 500x, to expand this dataset into a pool of 14,175 different conditions. Depth ratios were calculated accordingly, gnomAD allele frequencies, segmental duplication, and repeat masker information was annotated.

The entire BioData1 and random subsampling from SimData1 were combined to generate a training and validation dataset with approximately 200,000 variants from the 1,000,000 training variants. 180,000 variants were selected for model training, 45% from SimData1 and 55% from resampling of BioData1. This dataset was used for the model training and evaluation of the sensitivity and specificity of the selected model, and their features including AF distribution and biological appearances were very similar to published biological data (Supplementary Fig. 1).

#### BioData2

To estimate the performance of the pre-trained models and select the model with the best performance for DeepMosaic-CM, we introduced an independent gold-standard dataset. Variants were computationally detected from replicated sequencing experiments generated from 6 distinct sequencing centers and validated in 5 different centers, known as the common reference tissue project from the Brain Somatic Mosaicism Network^23^. 400 variants underwent multiple levels of computational validations including haplotype phasing, CNV exclusion, population shared exclusion, as well as experimental validation such as whole-genome single cell sequencing, Chromium Linked-read sequencing (10X Genomics), PCR amplicon sequencing, and droplet digital PCR. After validation, 43 true positive MVs and 357 false positive variants were determined as gold-standard evaluation set for low-fraction single nucleotide MVs from the 250x WGS data^23^. We extracted deep whole-genome sequences for those variants, labeled them accordingly and used them as gold standard validation set for model selection (Supplementary Fig. 2).

#### BioData3

To evaluate the performance of DeepMosaic-CM trained on a different portion of biological variants, we included another large-scale validation experiment we recently generated. Variant information and raw sequencing reads of 300x PCR-free PE150-only WGS of 18 samples from 9 different brain regions, cerebellum, heart, liver, and both kidneys of one individual were extracted from the capstone project of the Brain Somatic Mosaicism Network^18^. 1400 genomic positions with variants identified from WGS sample and reference homozygous/heterozygous controls validated by orthogonal experiments were selected and labeled as positive and negative according to the experimental validation result. The 1400 genomic positions were also down- and up-sampled for a targeted depth of 30, 50, 100, 150, 200, 250, 300, 400, and 500x. Depth ratios were calculated accordingly, gnomAD allele frequencies, segmental duplication, and repeat masker information were annotated.

#### SimData2

To compare the performance of DeepMosaic and other software to detect mosaicism on simulated data, we randomly generated another simulation dataset, with the following modifications: 1] only 7610 variants on non-repetitive region of chromosome 22 were considered true positive genomic positions; 2] random errors were generated from the Illumina NovaSeq sequencer error model. 3] Data was randomly down-sampled and up-sampled for a targeted depth of 50, 100, 200, 300, 400, and 500x. A total of 439,200 different variants were generated. FASTQ files were aligned and processed with BWA (v0.7.17), SAMtools (v1.9), and Picard (v2.18.27). The data was subjected to DeepMosaic as well as MuTect2 (GATK v4.0.4, both paired mode and single mode), Strelka2 (v2.9.2), MosaicHunter (v1.0.0), and MosaicForecast (v8-13-2019) with different models trained for different read depth (250x model for depth≥300x).

#### SimData3

We further generated another simulation dataset in a way that was fundamentally different from the training data with a positive:negative ratio similar to real data^18^ to compare the performance of DeepMosaic and other software for the detection of mosaic variants. We selected 30,090 genomic positions with reference homozygous genotype from a different genomic region (the entire Chromosome 1) of the whole-genome deep sequences from the ‘Genome In a Bottle’ sample HG002 (NA24345)^26^. The genomic positions from the 30,090 positions were genotyped as homozygous and fulfilled additional criteria 1] zero alternative bases in the raw sequencing data; 2] no detectable insertions/deletions in the position of interest; 3] have a genomic distance of at least 1000 bases between each other. On this clear background, 15,471 of them were labeled as “true negative” with reference homozygous genotype, 6868 were labeled as “true positive” mosaic variants with expected alternative AF 0.01, 0.02, 0.03, 0.04, 0.05, 0.10, 0.15, 0.20, and 0.25 (on average 763 variants for each genotype); 7751 were labeled as “true negative” heterozygous variants with alternative AF 0.50; the latest version of a different software BAMSurgeon (updated 24 Dec 2020) was used to generate this simulation dataset and retain the sequencing errors from the original biological samples. The original bam file was first up-sampled, alternative reads were replaced to generate the expected AF, mapped back to the genome and merged back to the bam file, according to the software manual^25^. Bam files with and without simulated data were downsampled to 500x, 400x, 300x, 200x, 100x, and 50x. The data were subjected to DeepMosaic as well as MuTect2 (GATK v4.0.4, both paired mode and single mode), Strelka2 (v2.9.2), MosaicHunter (v1.0.0), and MosaicForecast (v8-13-2019) with different models trained for different read depth (250x model for depth≥300x), the performance of the 180,540 points were evaluated.

#### BioData4

This additional dataset was used to compare the performance of DeepMosaic and other mosaic variant callers on biological samples. 16 WGS samples from blood and sperm of 8 individuals were sequenced at 200x^27^ (PRJNA588332). WGS was performed using an Illumina TrueSeq PCR-free kit with 350bp insertion size and sequenced on an Illumina HiSeq sequencer. Reads were aligned to the GRCh37 genome with BWA (v0.7.15) *mem* and duplicates were removed with sambamba (v0.6.6) and base quality recalibrated by GATK (v3.5.0). Processed BAM files were subjected to DeepMosaic as well as MuTect2 (GATK v4.0.4, both paired mode and single mode), Strelka2 (v2.9.2), MosaicHunter (v1.0.0), and MosaicForecast (v8-13-2019) with 200x models trained for the specific depth. Data from one of the individuals (F02) was down-sampled to 150x, 100x, 50x, and 30x with the SAMtools (v1.9) *view* command for the further benchmark of DeepMosaic.

### Neural network building and model training

For the 10 neural network architectures, Inception-v3, Resnet and Densenet were imported from PyTorch’s (v1.4.0) built-in library, while the 7 different builds of EfficientNet were imported from the efficientnet_pytorch (v0.6.1) Python (v3.7.1) package. The final fully connected layer of each model was replaced to be fed into 3 output units representing intermediate results instead of the default 1,000 output units for the 1,000 ImageNet classes to significantly reduce the total images required to extract basic features such as edges, stripes from raw images. A transfer-learning method was adopted for model training. Each model’s initial pre-trained weights provided by Pytorch and efficientnet_pytorch packages were trained on the ImageNet dataset. Before model training, we randomly divided the entire training dataset (including down-sampling and up-sampling of SimData1 and BioData1) to 80% “training” and 20% “evaluation” sets and fixed the split during model training while shuffling the order within the training set and evaluation set for each training epoch to form mini-batches for gradient descent. Each network architecture was trained using a batch size of 20 with a stochastic gradient descent (SGD) optimizer with learning rate of 0.01, and momentum of 0.9. The training was terminated until the training losses plateaued and evaluation accuracy reached 90% for each model architecture. The training was conducted on NVIDIA Kepler K80 GPU Nodes on San Diego Supercomputer Centre’s Comet computational clusters.

### Network selection

To select the “best-performing” neural network architecture among the trained Inception-v3, Resnet, Densenet and 7 different builds of EfficientNet, the gold standard evaluation dataset (BioData2) was used to test each model’s performance on biological (non-simulated) MVs determined by the dataset. Accuracy, MCC, True positive rates were calculated for each model and in the end EfficientNet-b4 at epoch 6 with the highest Accuracy, MCC and True positive rate among all model architectures was selected as our DeepMosaic model. The performance of DeepMosaic model (EfficientNet-b4 architecture) was further evaluated.

### Independent model training and evaluation for DeepMosaic-CM

To evaluate the performance of DeepMosaic-CM when trained on a different portion of biological variants, 15 epochs were trained for the EfficientNet-b4 architecture on 5 different training sets consisting of 122, 424 genomic positions. EfficientNet was imported from the efficientnet_pytorch (v0.6.1) Python (v3.7.1) package. The 5 different training sets were generated based on SimData1, BioData1, SimData2, and BioData3. 1] BioData only: 40,808 variants from the entire of BioData1 and BioData3 were pooled. The overall positive:negative ratio was 26.8%:73.2%. 2] SimData only for SimData1: 40,808 variants were selected from SimData1 with the matched number of positive and negative labels as BioData only. 3] SimData only for SimData2: 40,808 variants that were agreed by both MuTect2 and Strelka2 as “positive” or agreed by both methods as “negative” were selected from SimData2 with the matched number of positive and negative labels compared to BioData only. 4] BioData+SimData for SimData1: 40,808 variants half from BioData half from SimData only for SimData1 were selected with the matched number of positive and negative labels compared to BioData only. 5] BioData+SimData for SimData2: 40,808 variants half from BioData half from SimData only for SimData2 were selected with the matched number of positive and negative labels compared to BioData only. Each network architecture was trained using a batch size of 4 with a stochastic gradient descent (SGD) optimizer with a learning rate of 0.01, and momentum of 0.9. Fifteen different epochs were trained on each of the 5 training sets described above, and the model after each epoch is saved for performance evaluation. The training was conducted on NVIDIA GTX 980 GPU Nodes on San Diego Supercomputer Center’s Triton Shared Computing Cluster (TSCC). The training performance of the models was further evaluated on BioData2, which has not been used for any of the training procedures.

### Usage of DeepMosaic

Detailed instructions for users as well as the demo input and output is provided on GitHub (https://github.com/Virginiaxu/DeepMosaic).

### Orthogonal validation with deep amplicon sequencing

Deep amplicon sequencing analysis was applied to 241 variants from the 1355 candidates detected by all 5 mosaic variant callers from the 200× WGS of 16 samples^27^ to experimentally confirm the validation rate of DeepMosaic as well as other methods. PCR products for sequencing were designed with a target length of 160-190 bp with primers being at least 60 bp from the base of interest. Primers were designed using the command-line tool of Primer3^33^ with a Python (v3.7.3) wrapper. PCR was performed according to standard procedures using GoTaq Colorless Master Mix (Promega, M7832) on sperm, blood, and an unrelated control. Amplicons were enzymatically cleaned with ExoI (NEB, M0293S) and SAP (NEB, M0371S) treatment. Following normalization with the Qubit HS Kit (ThermoFisher Scientific, Q33231), amplification products were processed according to the manufacturer’s protocol with AMPure XP Beads (Beckman Coulter, A63882) at a ratio of 1.2x. Library preparation was performed according to the manufacturer’s protocol using a Kapa Hyper Prep Kit (Kapa Biosystems, KK8501) and barcoded independently with unique dual indexes (IDT for Illumina, 20022370). The libraries were sequenced on a NovaSeq platform with 100 bp paired-end reads. Reads from deep amplicon sequencing were mapped to the GRCH37d5 reference genome by BWA mem and processed according to GATK (v3.8.2) best practices without removing PCR duplicates. Putative mosaic sites were retrieved using SAMtools (v1.9) mpileup and pileup filtering scripts described in previous TAS pipelines^27^. Variants were considered positively validated for mosaicism if 1] their lower 95% exact binomial CI boundary was above the upper 95% CI boundary of the control; 2] their AF was >0.5%. The number of reference and alternative alleles calculated from the Amplicon validation was provided in Supplementary Table 1.

### Analysis of different categories of variants overlap with different genomic features

In order to assess the distribution of MVs and their overlap with genomic features, an equal number of variants (mSNVs/INDELs as in group G1-G7 in Supplementary Fig. 6) was randomly generated with the BEDtools (v2.27.1) shuffle command within the region from Strelka2 without the subtracted regions (e.g. repeat regions). This process was repeated 10,000 times to generate a distribution and their 95% CI. Observed and randomly subsampled variants were annotated with whole-genome histone modifications data for H3k27ac, H3k27me3, H3k4me1, and H3k4me3 from ENCODE v3 downloaded from the UCSC genome browser (http://hgdownload.soe.ucsc.edu/goldenPath/hg19/database/)—specifically for the overlap with peaks called from the H1 human embryonic cell line (H1), as well as peaks merged from 10 different cell lines (Mrg; Gm12878, H1, Hmec, Hsmm, Huvec, K562, Nha, Nhek, and Nhlf). Gene region, intronic, and exonic regions from NCBI RefSeqGene (http://hgdownload.soe.ucsc.edu/goldenPath/hg19/database/refGene.txt.gz); 10 Topoisomerase 2A/2B (Top2a/b) sensitive regions from ChIP-seq data (Samples: GSM2635602, GSM2635603, GSM2635606, and GSM2635607); CpG islands: data from the UCSC genome browser (http://hgdownload.soe.ucsc.edu/goldenPath/hg19/database/); genomic regions with annotated early and late replication timing^34^; high nucleosome occupancy tendency (>0.7 as defined in the source, all values were extracted and merged) from GM12878; enhancer genomic regions from the VISTA Enhancer Browser (https://enhancer.lbl.gov/); and DNase I hypersensitive regions and transcription factor binding sites from Encode v3 tracks from the UCSC genome browser (wgEncodeRegDnaseClusteredV3 and wgEncodeRegTfbsClusteredV3, respectively).

## Supplementary Information

**Supplementary Figure 1.**
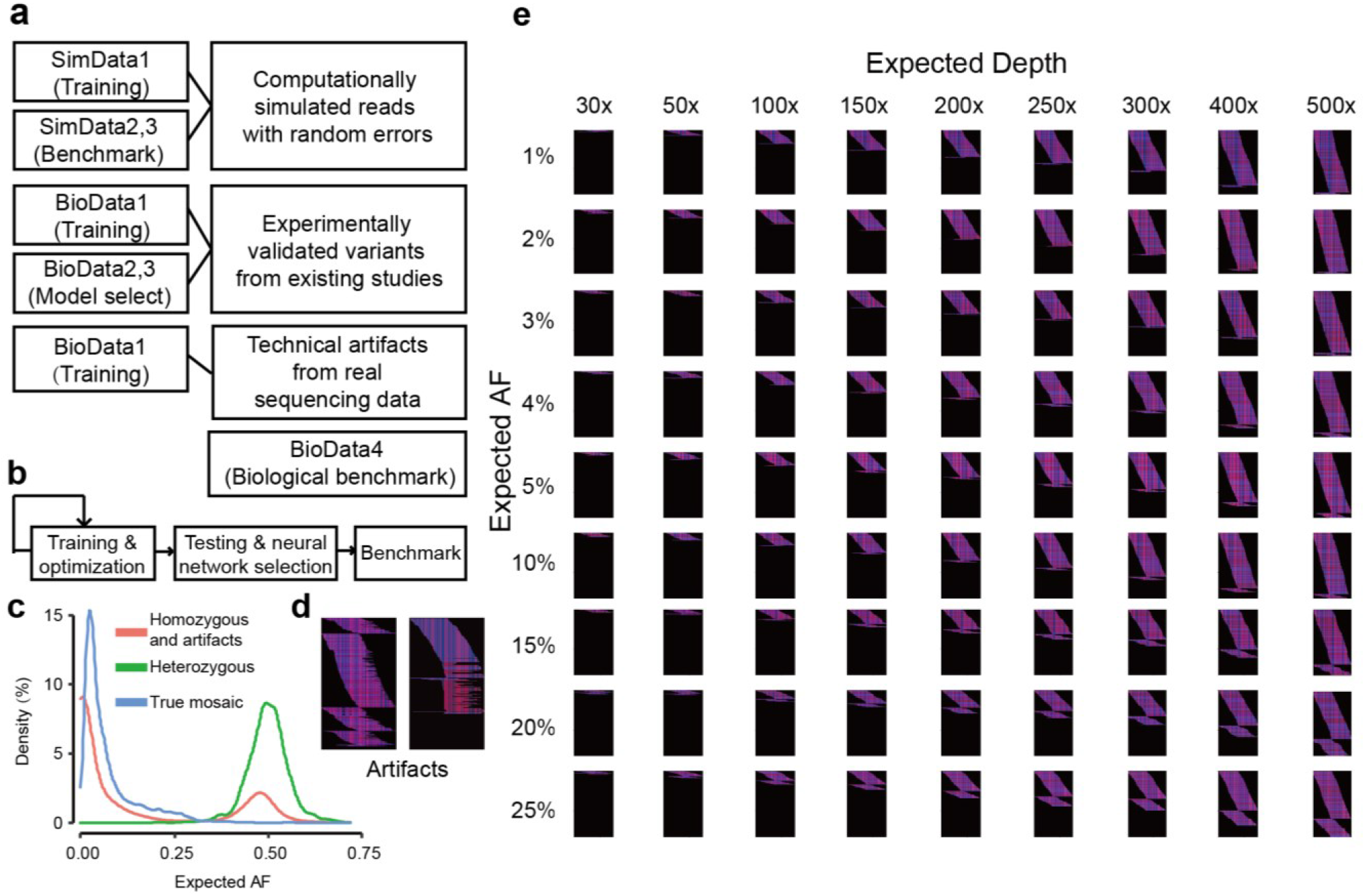
Training strategies and examples of training data for DeepMosaic. **(a)** More than 200,000 training and validation variants were generated for DeepMosaic, including computational simulations (SimData1), biologically validated variants from existing studies with manually curated technical artifacts (BioData1). We further included 1 gold standard dataset for testing and model selection (BioData2); all selected positive or negative variants underwent amplicon sequencing in at least one tissue sample according to the publication. We further included independent simulated data (SimData2 and SimData3) and biological data (BioData3 and BioData4) to benchmark DeepMosaic. **(b)** The overall strategies of model training and benchmarking for each tested model. **(c)** The distribution of probability density of expected AFs for different variants from the training set. Red: Reference homozygous variants and technical artifacts are labeled “Negative” in the training set. Green: Heterozygous variants are also labeled “Negative” in the training set. Blue: True mosaic variants are labeled “Positive” in the training set. **(d)** Two examples of false positive variants with different sequencing artifacts, left: multiple alternative alleles from sequencing bias or alignment artifacts; right: reads truncated because of sequencing or alignment artifacts. **(e)** All training images were down-sampled and up-sampled into 30x, 50x, 100x, 150x, 200x, 250x, 300x, 400x and 500x, mutant allelic fractions (AFs) from the simulated data that were set as 1%, 2%, 3%, 4%, 5%, 10%, 15%, 20%, 25% and shown.

**Supplementary Figure 2.**
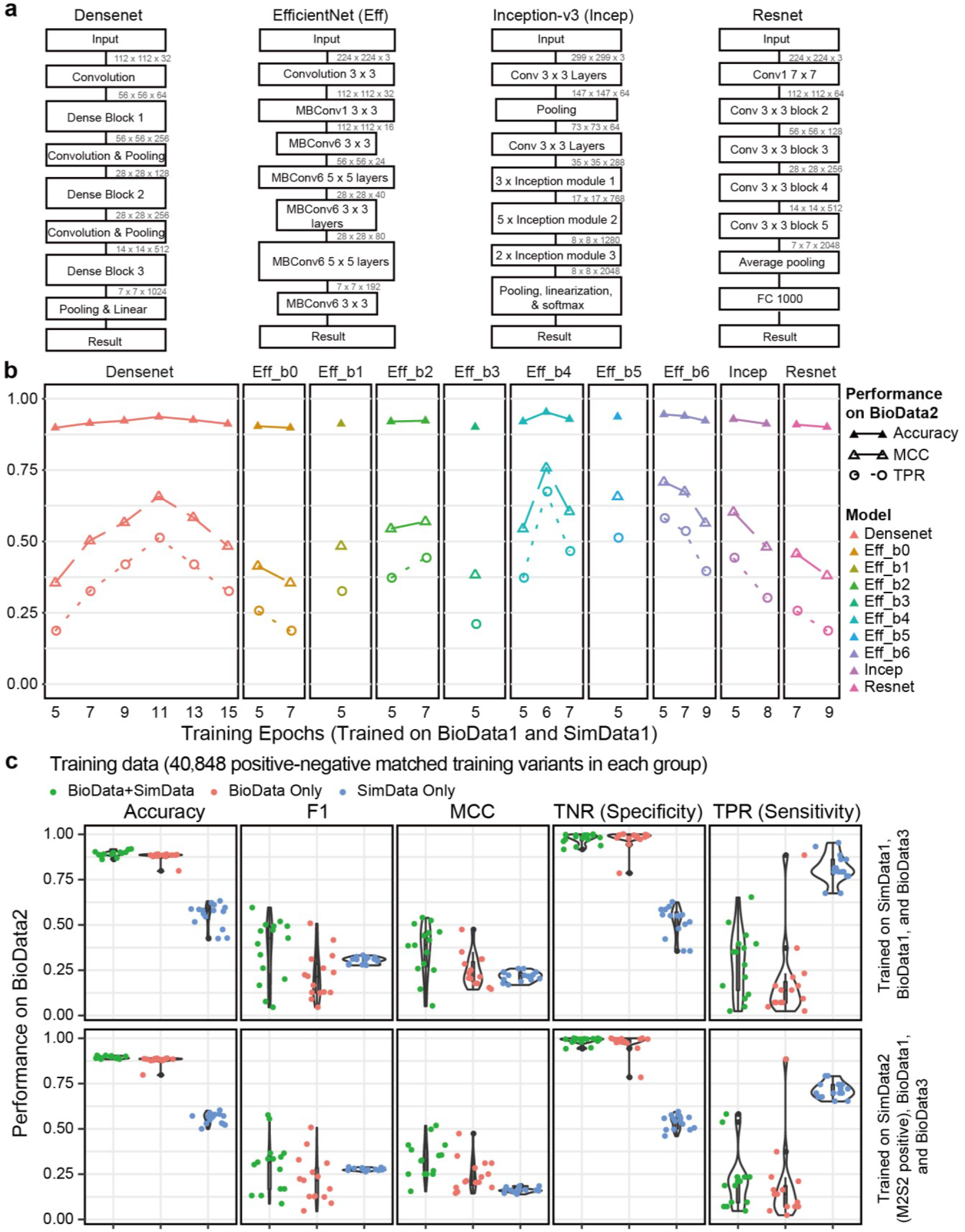
Network model selection based on independent gold-standard testing set. **(a)** Comparison of network structures implementing a variety of classification algorithms. For different build versions of EfficientNet, only a general structure is shown. Inception v3 was used in DeepVariant, and Resnet was used in NeuSomatic. **(b)** All models were trained on 180,000 training variants from BioData1 and SimData1 until the models reach training accuracy > 0.9. Accuracy, Matthews’s correlation coefficient (MCC), and Sensitivity of different network structures trained with the same data with different epochs. EfficientNet-b4 trained at 6 epochs demonstrated the highest Accuracy, MCC, and Sensitivity on the gold standard validation set^23^ (BioData2); thus it was used as the default core model for DeepMosaic. We additionally provide an option for experienced users to train their own models with self-labeled training data. **(c)** EfficientNet-b4 models were trained on 5 additional datasets, each for 15 epochs. The training datasets were generated with different compositions of biologically validated data and simulated data. Models trained only on simulated data showed overall higher sensitivity but much lower specificity on the gold standard evaluation set (BioData2) due to the high fraction of false-positive calls. Models trained only on biological data showed similar overall performance compared with models trained on a mixture of biological and simulated data. All three training sets are generated with the same number of positive and negative data points as the biological data and with the same number of total variants. M2S2 Positive: training variants were labeled positive by both MuTect2 and Strelka2.

**Supplementary Figure 3.**
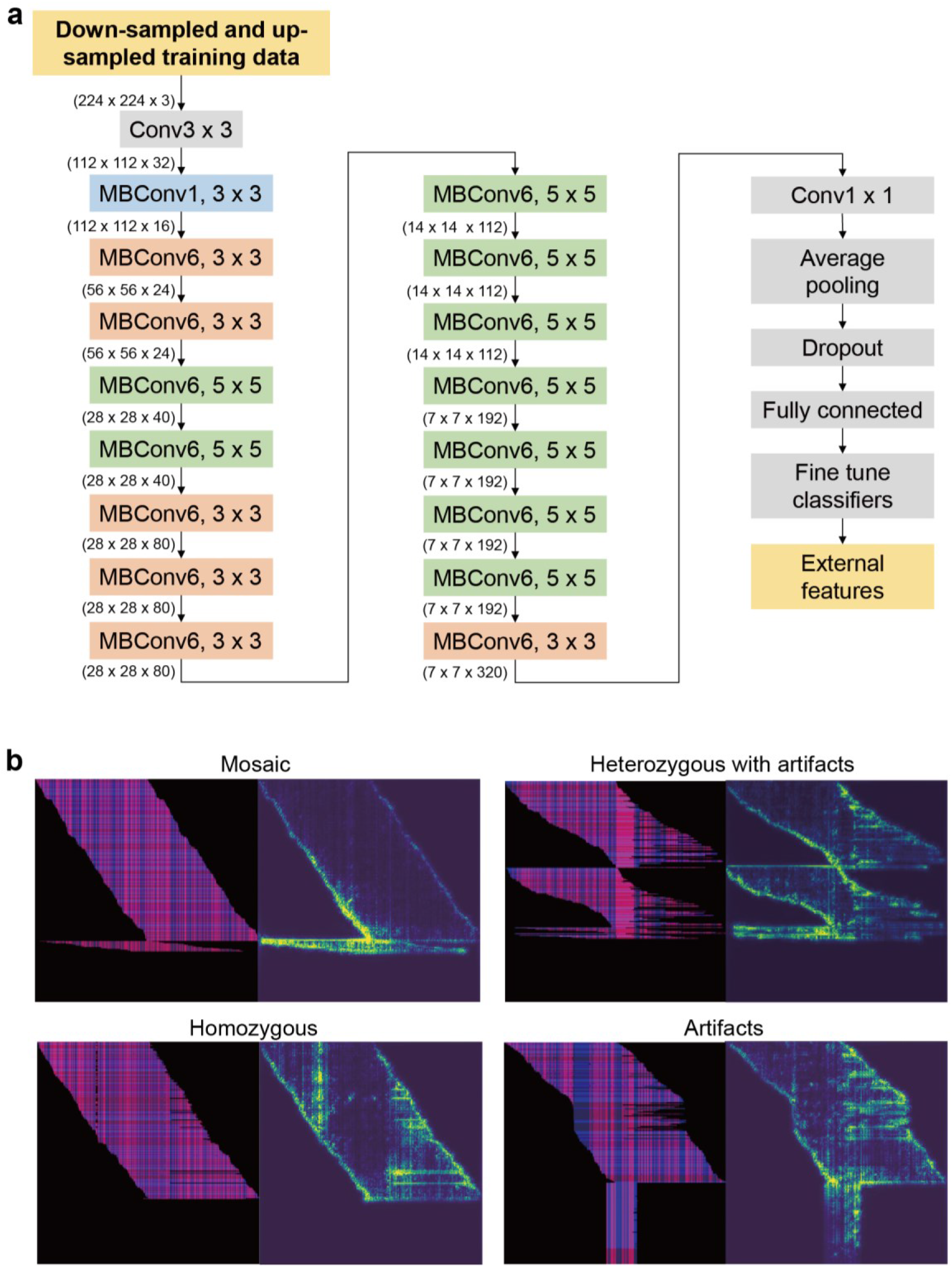
The convolutional neural network of the DeepMosaic default model and gradient visualization with guided backpropagation for the DeepMosaic default model (EfficientNet-b4). **(a)** Down- sampled and up-sampled image files coded from original BAM files were used as input. 16 mobile convolutional layers were adapted from EfficientNet-b4, with optimized parameter size and structures. Numbers represent the dimensions of trained hyperparameters. **(b)** A mosaic, a homozygous, and a heterozygous variant with artifacts, as well as a technical artifact, are shown here for the gradient visualization with guided backpropagation method23 implemented for the DeepMosaic core model, EfficientNet-b4 trained at epoch 6, left: image coding, right: gradient heatmap. The edges of bases, the sequence information, as well as other high-dimensional information, are highlighted by the model.

**Supplementary Figure 4.**
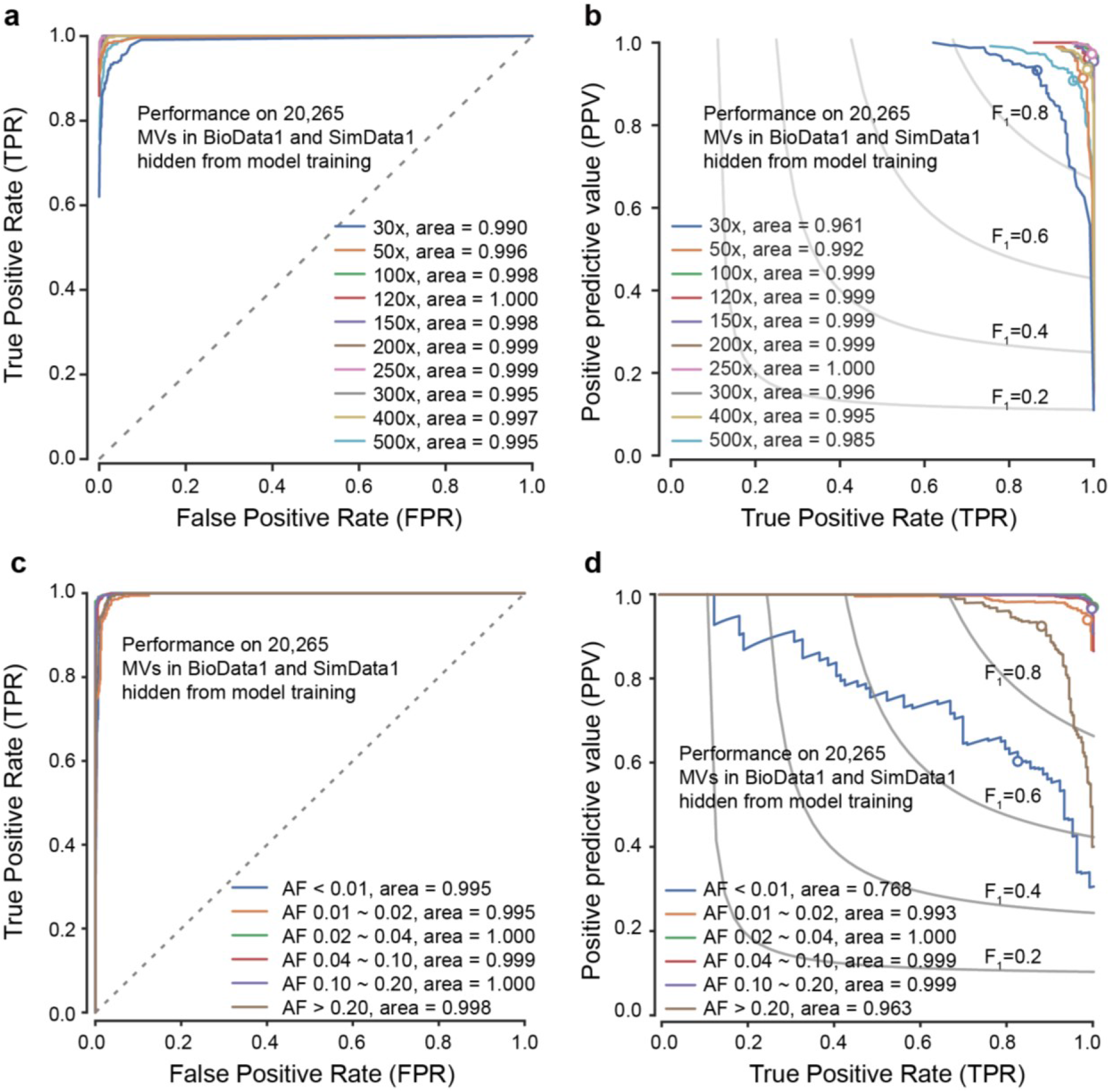
Performance of DeepMosaic default model (EfficientNet-b4) on data hidden from training. **(a)** Receiver operating characteristic (ROC) curve for DeepMosaic. True positive rates (TPR) and false-positive rates (FPR) were evaluated from 20,265 variants (BioData1 and SimData1) hidden from model training and model selection. Colors show groups of intended read depth. **(b)** Precision-recall curves for DeepMosaic, evaluated from the 20,265 hidden variants, dots showed the performance of the default parameters for DeepMosaic-CM. **(c)** ROC curve for DeepMosaic. TPR and FPR were evaluated from 20,265 variants (BioData1 and SimData1) hidden from model training and model selection. Colors show groups of bins of different expected AFs. **(d)** Precision-recall curves for DeepMosaic, evaluated from the 20,265 hidden variants, dots showed the performance of the default parameters for DeepMosaic-CM for different AF bins. Iso-F1 curves were shown for each precision-recall pairs with identical F1 scores labeled in (b) and (d).

**Supplementary Figure 5.**
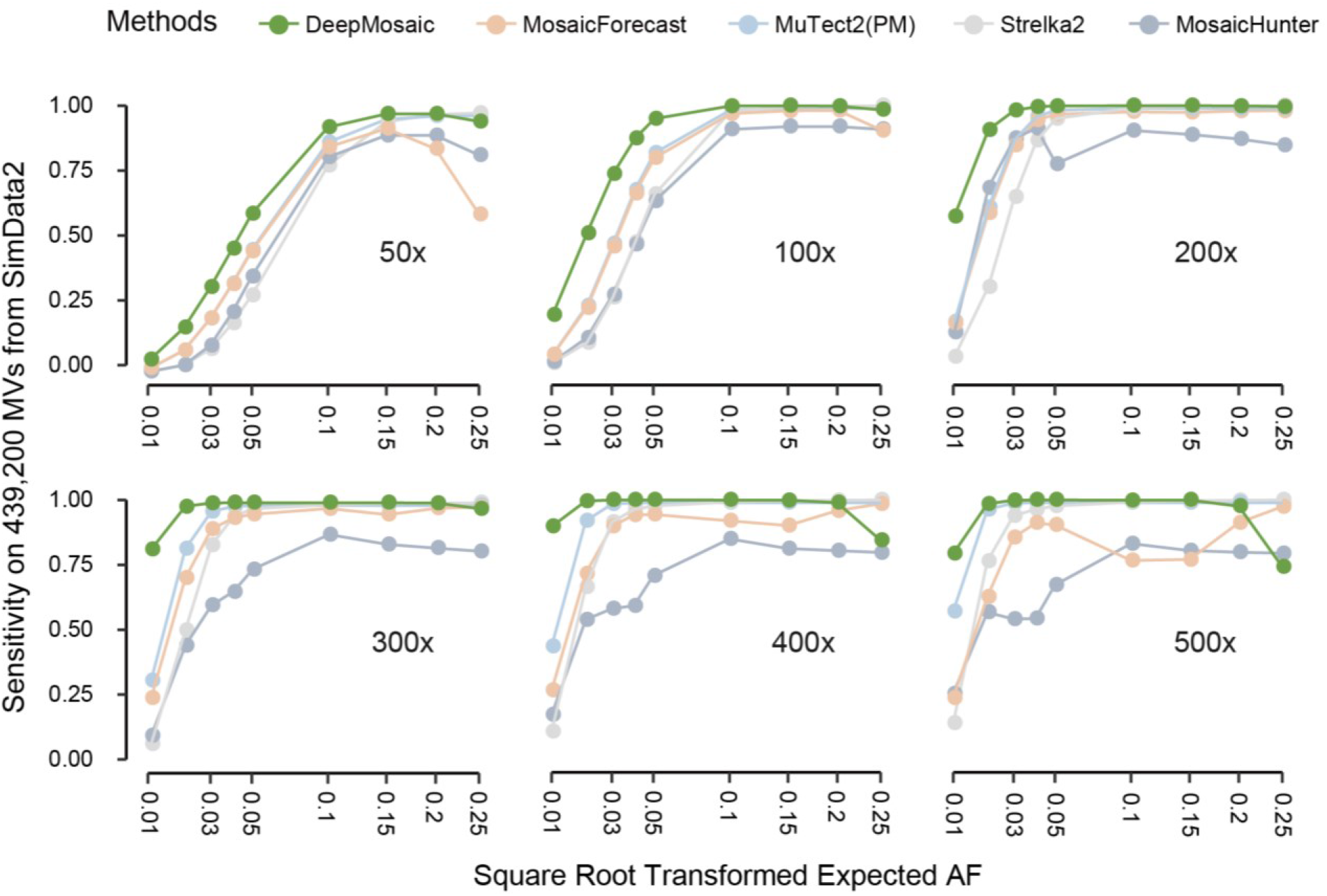
Performance of DeepMosaic and other mosaic variant callers on SimData2. Sensitivity of DeepMosaic and other mosaic callers on 439,200 independently simulated benchmark variants (SimData2) at simulated read depths and AFs. DeepMosaic performed equally well or better than other tested methods, especially at lower expected AFs.

**Supplementary Figure 6.**
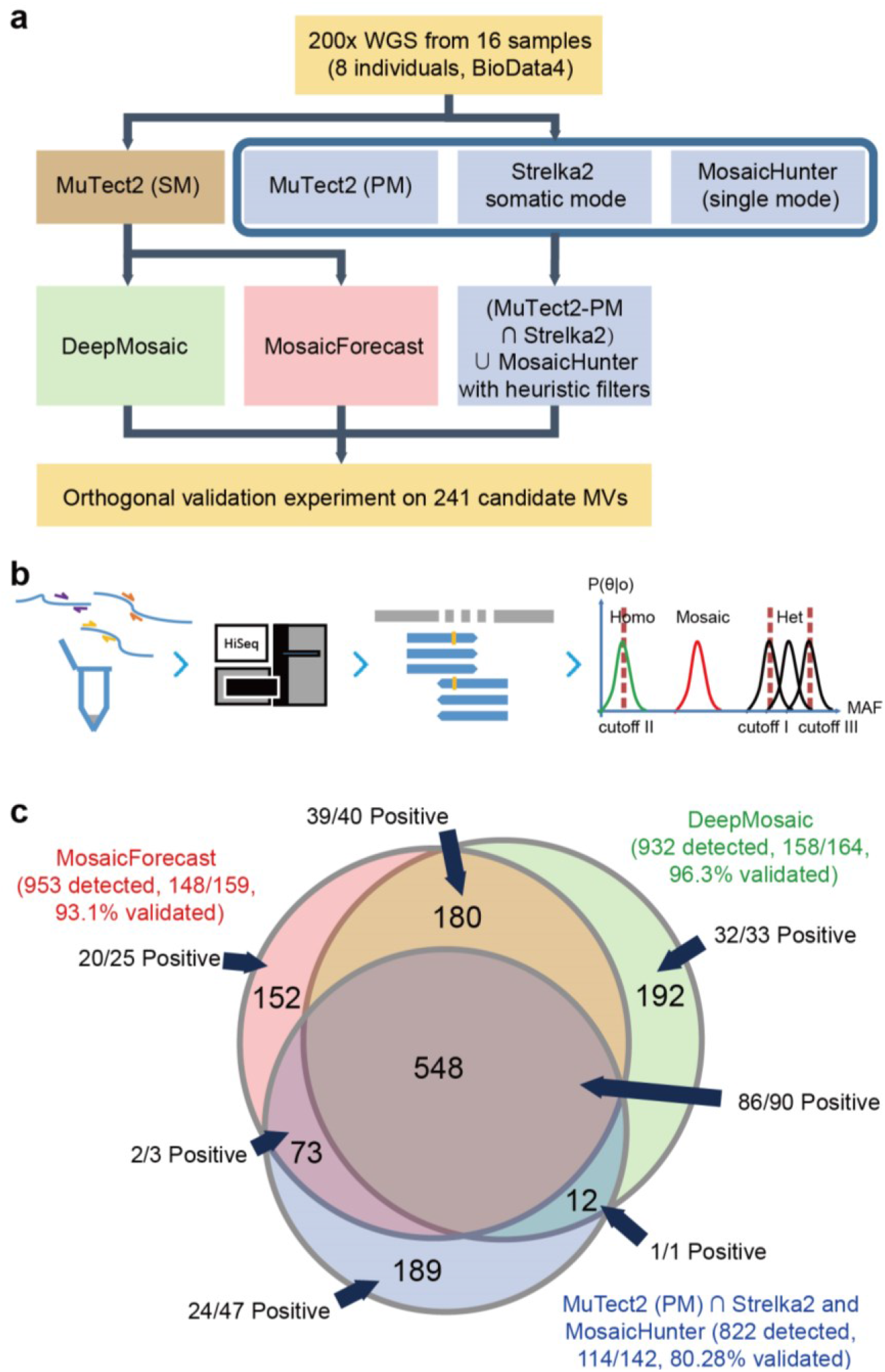
Comparison of DeepMosaic and traditional mosaic variant calling strategies on a biological dataset (BioData4) **(a)** Compared with the mosaic variant calling strategy (M2S2MH) used in a previous publication, DeepMosaic and MosaicForecast strategies are also listed. **(b)** Schematics for amplicon validation. Primers were designed for different candidates and amplicons were collected for Illumina sequencing. Information from aligned reads were calculated and genotypes were determined. **(c)** Venn diagram of the experimentally validated results and the portions of variants from different study strategies. DeepMosaic demonstrated a 96.3% (158/164) validation rate. Of all the 932 variants identified by DeepMosaic, 39.91% (372/932) were missed by the MuTect2 Strelka2 MosaicHunter pipeline^27^ with validation rate 97.26 (71/73) and 21.89% (204/932) were missed by the MosaicForecast^4^ pipeline with validation rate 97.06 (33/34).

**Supplementary Figure 7.**
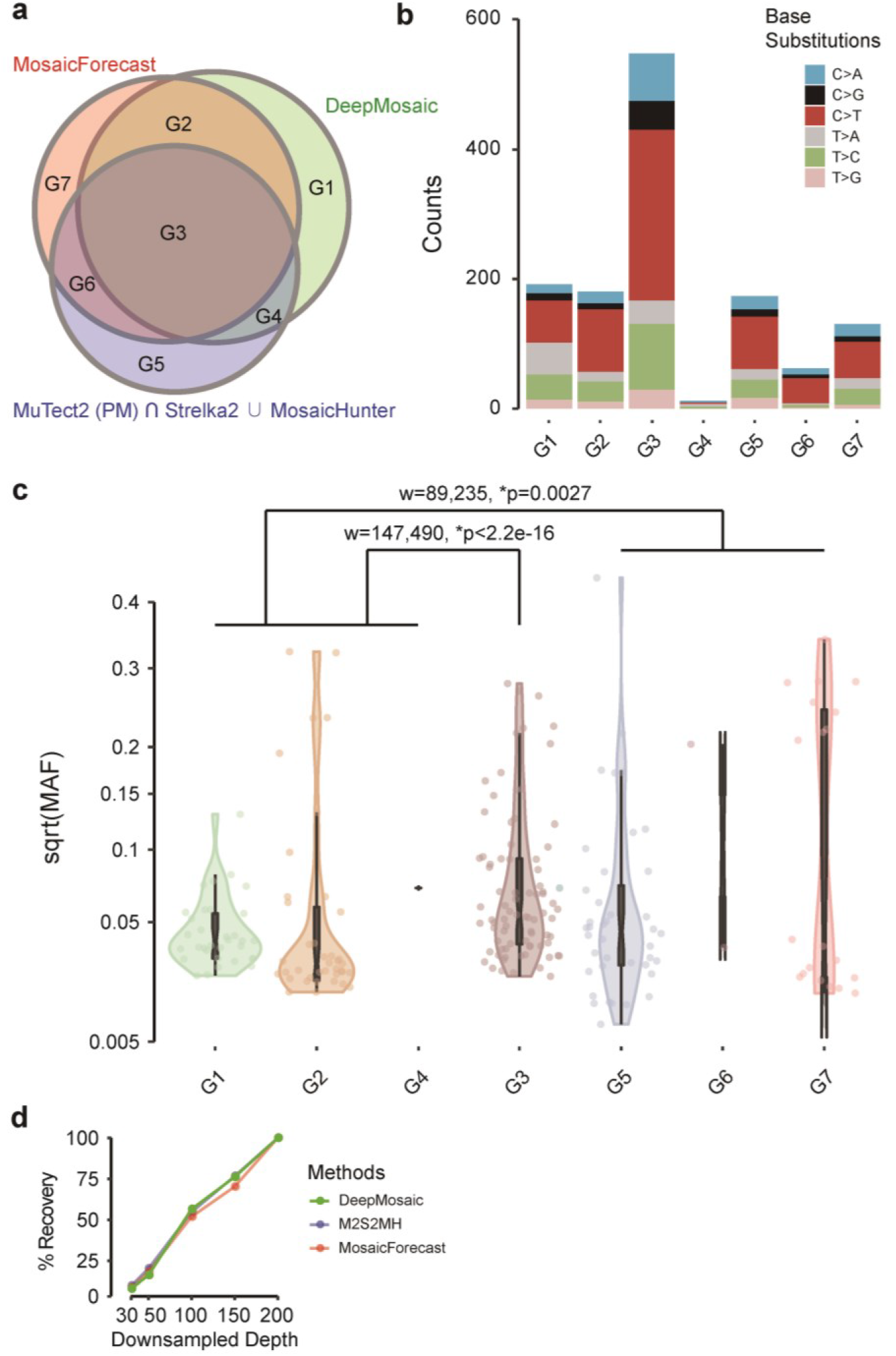
Comparison of features of variants called by DeepMosaic and other pipelines. **(a)** Different overlapping groups of variants detected by the 3 pipelines were separated into 7 groups. **(b)** DeepMosaic-specific (G1) variants present similar base-substitution features compared with variants detected by the MuTect2-Strelka2-MosaicHunter combined pipeline as well as MosaicForecast pipeline (G2-G7). **(c)** Allelic fractions of the variants detected in the original WGS sample showed that DeepMosaic-specific variants (G1, G2, and G4) showed a significantly lower average AF than variants detectable by all 3 pipelines (G3, p<2.2e-16 by a two-tailed Wilcoxon rank sum test with continuity correction) and lower than variants detectable only in other pipelines (G5, G6, and G7, p=0.0027 by a two-tailed Wilcoxon rank sum test with continuity correction). **(d)** Recovery rate of DeepMosaic, M2S2MH, and MosaicForecast at different depths from down sampling of BioData3. DeepMosaic showed a similar variant recovery rate compared with M2S2MH and MosaicForecast, even when considering the lower AF variants detected by DeepMosaic.

**Supplementary Figure 8.**
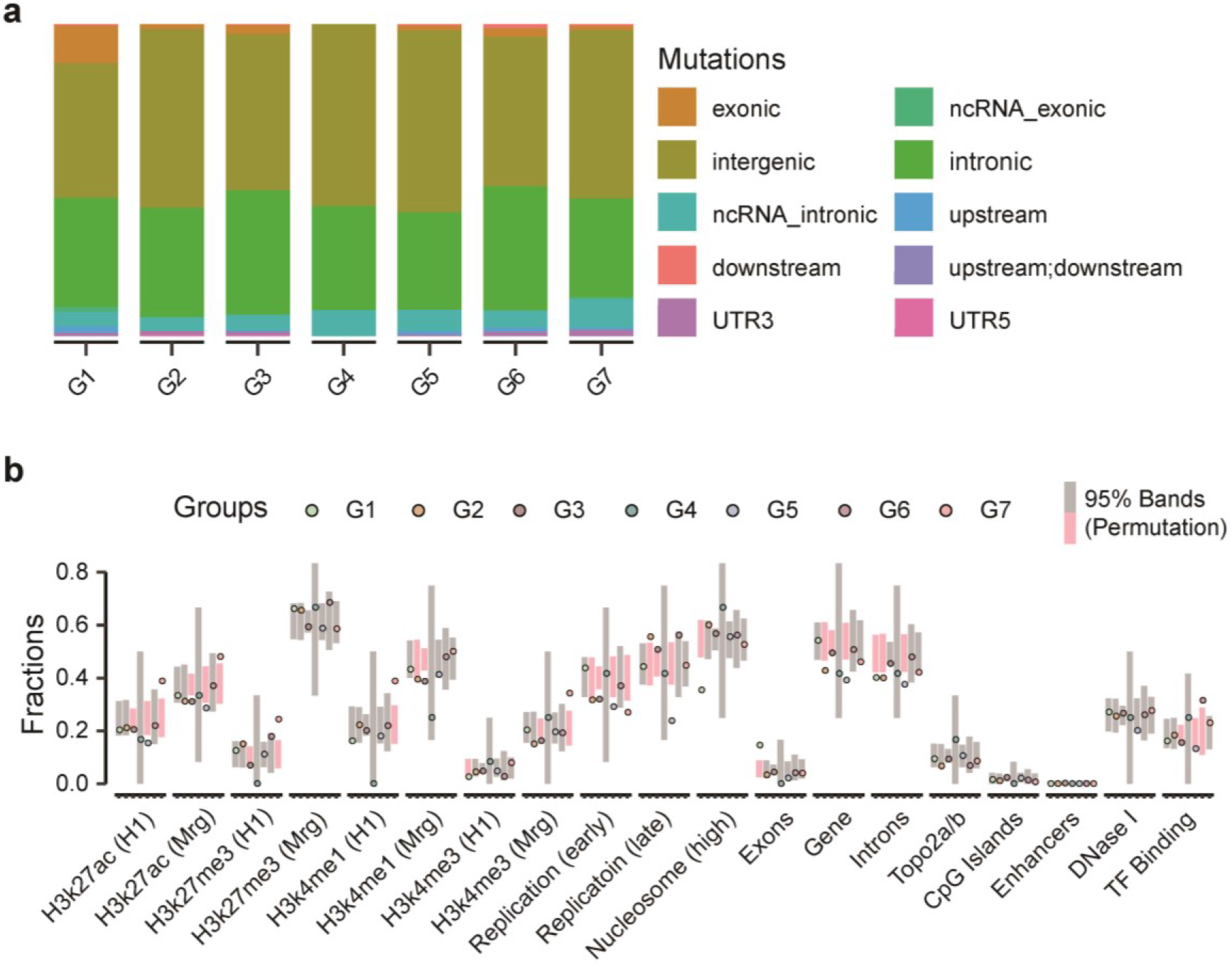
Enrichment of genomic features for variants called by DeepMosaic and conventional methods. **(a)** Variants called from different pipelines shared similar variant types and contributions. The groups are defined the same as Supplementary Fig. 6a. The relative contribution of different types of MVs is stable between different variant groups. **(b)** Enrichment analysis of variants in different genomic features. Unlike the variants shared with other callers, DeepMosaic-specific (G1) variants present depletion in high nucleosome occupancy regions. 10,000 permutation was carried out on randomly selected gnomAD variants, significant comparisons are shown in pink. Overall DeepMosaic-specific variants (G1) do not show significantly different genomic features compared with permutation intervals.

## Supplementary Text: Members of the Brain Somatic Mosaicism Network

Boston Children’s Hospital: August Yue Huang, Alissa D’Gama, Caroline Dias, Christopher A. Walsh, Javier Ganz, Michael Lodato, Michael Miller, Pengpeng Li, Rachel Rodin, Robert Hill, Sara Bizzotto, Sattar Khoshkhoo, Zinan Zhou

Harvard University: Alice Lee, Alison Barton, Alon Galor, Chong Chu, Craig Bohrson, Doga Gulhan, Eduardo Maury, Elaine Lim, Euncheon Lim, Giorgio Melloni, Isidro Cortes, Jake Lee, Joe Luquette, Lixing Yang, Maxwell Sherman, Michael Coulter, Minseok Kwon, Peter J. Park, Rebeca Borges-Monroy, Semin Lee, Sonia Kim, Soo Lee, Vinary Viswanadham, Yanmei Dou Icahn School of Medicine at Mt. Sinai: Andrew J. Chess, Attila Jones, Chaggai Rosenbluh, Schahram Akbarian

Kennedy Krieger Institute: Ben Langmead, Jeremy Thorpe, Jonathan Pevsner, Sean Cho

Lieber Institute for Brain Development: Andrew Jaffe, Apua Paquola, Daniel Weinberger, Jennifer Erwin, Jooheon Shin, Michael McConnell, Richard Straub, Rujuta Narurkar

Mayo Clinic: Alexej Abyzov, Taejeong Bae, Yeongjun Jang, Yifan Wang

Sage Bionetworks: Cindy Molitor, Mette Peters

Salk Institute for Biological Studies: Fred H. Gage, Meiyan Wang, Patrick Reed, Sara Linker

Stanford University: Alexander Urban, Bo Zhou, Xiaowei Zhu

Universitat Pompeu Fabra: Aitor Serres Amero, David Juan, Inna Povolotskaya, Irene Lobon, Manuel Solis Moruno, Raquel Garcia Perez, Tomas Marques-Bonet

University of Barcelona: Eduardo Soriano University of California, Los Angeles: Gary Mathern

University of California, San Diego: Danny Antaki, Dan Averbuj, Eric Courchesne, Joseph Gleeson, Laurel Ball, Martin Breuss, Subhojit Roy, Xiaoxu Yang

University of Michigan: Diane Flasch, Trenton Frisbie, Huira Kopera, Jeffrey Kidd, John Moldovan, John V. Moran, Kenneth Kwan, Ryan Mills, Sarah Emery, Weichen Zhou, Xuefang Zhao

University of Virginia: Aakrosh Ratan

Yale University: Alexandre Jourdon, Flora M. Vaccarino, Liana Fasching, Nenad Sestan, Sirisha Pochareddy, Soraya Scuderi

## Data availability

WGS data used to generate the training set are available at the Sequence Read Archive (SRA, Accession No. SRP028833 and SRP100797). The gold standard WGS data and validated capstone project data are available at the National Institute of Mental Health Data Archive (NIMH NDA Study ID 792 and 919: https://dx.doi.org/10.15154/1504248) and the Brain Somatic Mosaicism Consortium Data Portal. Simulated data generated from NA24385 (HG002) are available at https://humanpangenome.org/hg002/. The independent sperm and blood deep WGS data are available at SRA (Accession No. PRJNA588332).

## Code availability

DeepMosaic is implemented in Python; the code, documentation and demos are available at https://github.com/Virginiaxu/DeepMosaic.

## Acknowledgment

The authors thank Dr. Yanmei Dou for helping to set up the MosaicForecast pipeline. We thank Prof. Michael K. Gilson for the help in computational resources. We thank Profs. Peter J. Park, Garrison W. Cottrell, John V. Moran, Melissa Gymrek, Drs. Patrick J. Reed, August Y. Huang, and Si-Jin Cheng for their valuable comments and suggestions. This work was supported by the National Institute of Mental Health (U01MH108898), Rady Children’s Institute for Genomic Medicine and Howard Hughes Medical Institute. The authors thank San Diego Supercomputer Center (SDSC) (TG-IBN190021). This publication includes data generated at the UC San Diego IGM Genomics Center utilizing an Illumina NovaSeq 6000 that was purchased with funding from a National Institutes of Health SIG grant (#S10 OD026929).

## Contributions

X.Y., X.X., and J.G.G. conceived this project with input from M.B. and D.A.; X.Y. designed the study workflow and managed the project. X.X. implemented the image representation and neural network classifier under supervision and instruction by X.Y.; X.Y., C.L. and X.X. generated the training data with the help from D.A., R.D.G., and L.W.; X.X. performed the training and model selection under supervision by X.Y.; independent dataset were processed by M.B., D.A., and R.D.G. under supervision by J.S. and J.G.G.; X.Y. and M.B. performed the validation experiments with help from L.L.B. and C.C.; X.Y. and X.X. wrote the original and revised manuscript with input from all listed authors; X.Y. and J.G.G. edited the manuscript. DeepMosaic is benchmarked on part of the Brain Somatic Mosaicism Network (BSMN) common brain experiment and common analysis pipeline for SNVs contributed by Y.W., T.B. under supervision by A.A. and the BSMN capstone project contributed by M.B., X.Y., D.A., and X.X. under supervision by J.G.G.; J.G.G. supervised this project. All authors discussed the results and contributed to the final manuscript.

